# Alpha oscillations do not implement gain control in early visual cortex but rather gating in parieto-occipital regions

**DOI:** 10.1101/2020.04.03.021485

**Authors:** Alexander Zhigalov, Ole Jensen

## Abstract

Spatial attention provides a mechanism for respectively enhancing relevant and suppressing irrelevant information. While it is well-established that attention modulates oscillations in the alpha band, it remains unclear if alpha oscillations are involved in directly modulating the neuronal excitability associated with the allocation of spatial attention. In this study in humans, we utilized a novel broadband frequency (60 – 70 Hz) tagging paradigm to quantify neuronal excitability in relation to alpha oscillations in a spatial attention paradigm. We used magnetoencephalography to characterize ongoing brain activity as it allows for localizing the sources of both the alpha and frequency tagging responses. We found that attentional modulation of alpha power and the frequency tagging response are uncorrelated over trials. Importantly, the neuronal sources of the tagging response were localized in early visual cortex (V1) whereas the sources of the alpha activity were identified around parieto-occipital sulcus. Moreover, we found that attention did not modulate the latency of the frequency tagged responses. Our findings point to alpha band oscillations serving a downstream gating role rather than implementing gain control of excitability in early visual regions.

**Significance Statement:** By combining magnetoencephalography and a novel broadband frequency tagging approach, we show that spatial attention differently modulates alpha oscillations and neuronal excitability. Importantly, the sources of the alpha oscillations and tagging responses were spatially distinct and the alpha power and tagging response were not related over trials. These results are inconsistent with previous ideas suggesting that alpha oscillations are involved in gain control of early sensory regions; rather alpha oscillations are involved in the allocation of neuronal resources in downstream regions.

## Introduction

Attention provides a mechanism that allows enhancing relevant and suppressing irrelevant information (Kastner and Nobre, 2014). When several complex stimuli are presented in a visuo-spatial scene a selection mechanism is enhancing and suppressing relevant and irrelevant information, respectively. Attention results in competitive interactions among neurons, causing them to respond stronger to attended stimuli while distracting stimuli are suppressed (Desimone and Duncan, 1995). The underlying neuronal mechanisms of these modulations in relation to oscillatory brain activity remain elusive. Several studies showed that the power of alpha band oscillations is modulated by attention (Foxe and Snyder, 2011; Klimesch, 2012; Jensen and Hanslmayr, 2020) and it has been suggested that alpha oscillations modulates gain control mechanism by influencing neuronal excitability (Romei et al., 2008; Jensen and Mazaheri, 2010; Mathewson et al., 2011; Keitel et al., 2019; van Diepen et al., 2019). It is also known that attention modulates power of neuronal response to flickering (or tagging) stimuli at higher frequencies (Gulbinaite et al., 2019). Interestingly, while alpha power decreases contra-laterally to attended stimulus (and increases ipsi-laterally), power of the high-frequency tagging response shows an opposite effect (e.g. (Gulbinaite et al., 2019; Zhigalov et al., 2019). There have been some attempts to assess the relationship between power of the alpha oscillations and the tagging response in attentional tasks. These studies indicate that the frequency tagging signal is not related to alpha magnitude on a trial-by-trial basis (Zhigalov et al., 2019; Gundlach et al., 2020). This then begs the question of what is the functional role of alpha oscillations in attention tasks? Magnetoencephalography (MEG) offers the opportunity to localize the sources of the alpha oscillations and the frequency tagging response. If the sources are at different levels of the visual hierarchy, this would provide important clues to the functional role of alpha oscillations.

Broad-band frequency tagging also allows us to assess the delay of the neuronal response with respect to visual input. While it is well established that spatial attention can increase neuronal responses, it might also speed up processing, i.e. neurons in visual cortex responding to attended objects will fire earlier compared to those for unattended objects. A study by (Sundberg et al., 2012) showed that attention does produce small (1–2 ms) but significant reductions in the latency of both the spiking and LFP responses in extrastriate cortex V4. Similarly, another study demonstrated that both latency and peak amplitude of the response are modulated by attention, but only latencies correlate with reaction time (Galashan et al., 2013). These findings motivate exploring if response latencies change with attention in humans. As we will explain, processing delays can be estimated by cross-correlating the broad-band visual input with the MEG response.

In this study, we used a novel experimental design to dissociate the effect of attention on the alpha and tagging responses. We utilized a spatial attention task where the luminance of visual stimuli in the left and right visual field was driven by independent random broadband signals (60 – 70 Hz). This broadband frequency tagging technique in combination with magnetoencephalography (MEG) allowed us to obtain reliable responses to invisible stimulation that did not perturb the alpha oscillations, and also estimate response latencies by cross correlating the tagging signal and the brain response as a function of attention. This allows us to correlate the frequency tagged response and alpha power over trials. We applied source modelling to localize the neuronal activity associated with the alpha and tagging responses.

## Materials and Methods

### Participants

Twenty-four right-handed participants (14 females) with no history of neurological disorders partook in the study. Six of the subjects were excluded from the analysis: two of the subjects showed an excessive amount of eye blinks and saccades occurring in more than 80% of trials. Other four subjects showed a weak neuronal response (*p* > 0.05, t-test, compared to the baseline) to the broadband tagging signal which did not allow us to reliably estimate the latencies of the tagging response. It should be noted that the neuronal response to flicker is highly individual (Herbst et al., 2013). This left eighteen subjects for further analysis.

The study was approved by the local ethics committee (University of Birmingham, UK) and written informed consent was acquired before enrolment in the study. All subjects conformed to standard inclusion criteria for MEG experiments. Subjects had normal or corrected-to-normal vision. Subjects received financial compensation of £15 per hour or were compensated in course credits.

### Experimental paradigm

Participants performed a spatial attention task (8 blocks of 6 min) in which they were instructed to allocate attention to either the left or right visual hemifield in accordance to the cue presented at the beginning of each trial (Fig. 1).

**Figure 1.**
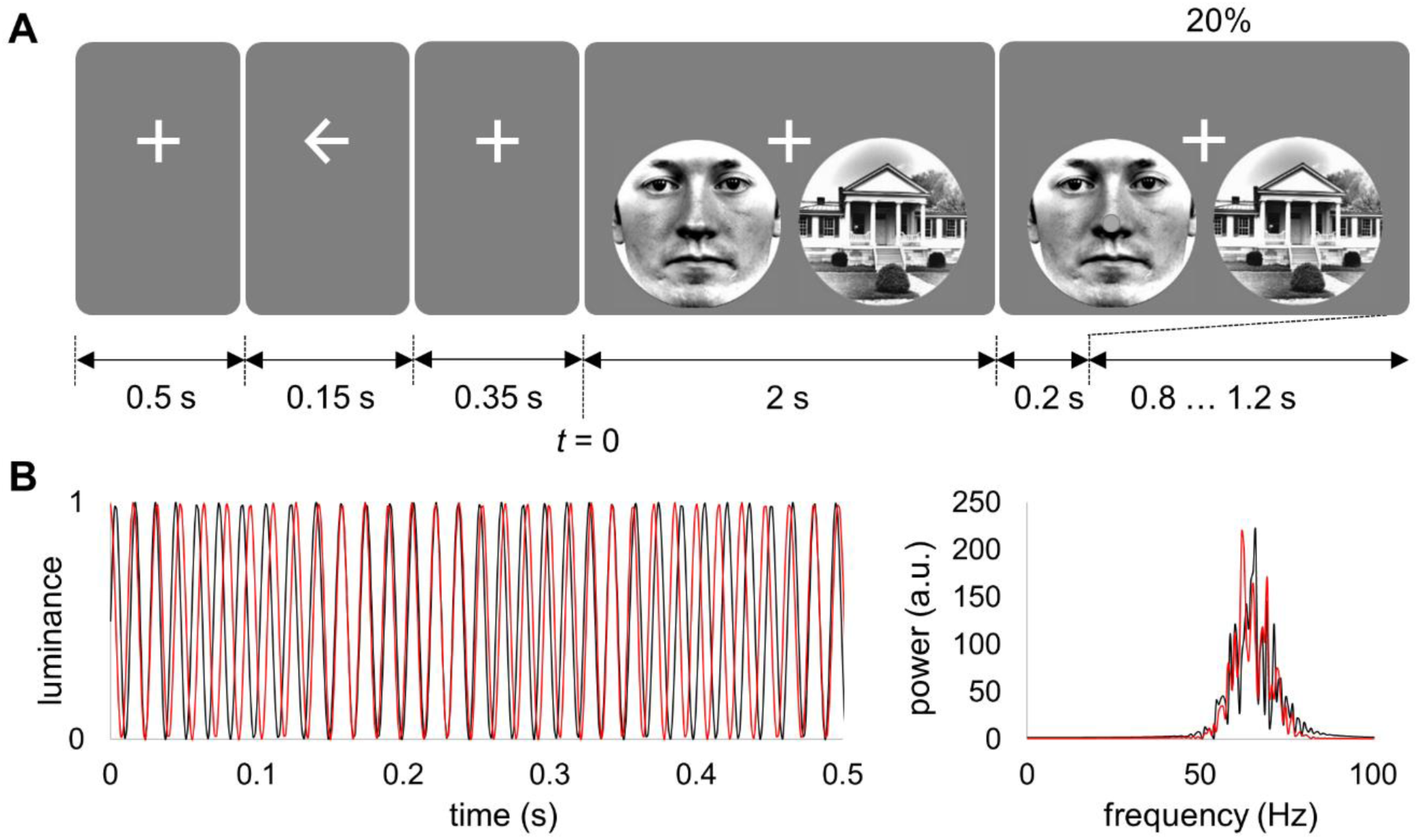
(A) The experimental paradigm. After a cue (left/right arrow), a house-face pair was presented superimposed with broadband (60 – 70 Hz) flicker signals. In 20% of the trials, a small part (1.2°) of the image flipped and required participant’s response. In 5% of the trials (catch trials), the flip was in the hemifield opposite to the cued side and participants had to ignore this event. The onset of frequency tagging is marked as *t* = 0. (B) Example of the two uncorrelated broadband signals (red and black lines depict left and right stimuli, respectively) used to drive the visual stimuli and the respective power spectra. Note that the broadband signals were generated independently for each trial.

Each trial started with a fixation cross (500 ms) followed by a cue (150 ms; left/right arrow) indicating the hemifield that the participants had to attend to, while fixating on the center of the screen. A fixation cross was shown for 350 ms after the attentional cue, and then stimuli were presented in the left and right visual hemifield for 2000 ms. Participants were instructed to detect a flip occurring at the center of the cued stimulus. The flips occurred at the end of the trial in 25% of trials. In 20% of these trials, the flip was on the cued side, while in 5% of the trials (catch trials), the flip was in the hemi-field opposite to the cued side, and participants did not have to respond. Participants responded to the flip by pressing a button by either left or right index finger (counterbalanced over subjects). The duration of the flip was adjusted using QUEST adaptive staircase procedure (Watson and Pelli, 1983) to attain 80% correct responses. The initial duration of the flip was 10 ms and it varied between 2 and 30 ms during the session controlled by the QUEST procedure. The validity of the responses was indicated on the screen as correct (“CORRECT”), incorrect (“INCORRECT”), or missed (“MISS”) response. The next trial started after a random interval of 800 ± 200 ms. Such relatively short inter-stimulus interval may influence the neuronal responses in the subsequent trials; however, the random stimulus onset reduces this effect. The experimental paradigm was implemented in MATLAB 2018b (Mathworks Inc., Natrick, USA) using Psychophysics Toolbox 3.0.11 (Kleiner et al., 2007).

### Visual stimuli

Pairs of rounded stimuli (face and house) were presented simultaneously in the lower left and right visual field (5.7° eccentricity), and the target stimulus (i.e., flip) to which participants instructed to respond was 1.2° in diameter. Different combinations of faces and houses (comprising ten faces and ten houses) were presented in random order over the trials. The luminance of the grayscale stimuli was normalized using the SHINE Toolbox for MATLAB (Willenbockel et al., 2010) (see, Fig. 1).

The luminance of the stimuli (face and house) was modulated by broadband signals (60 – 70 Hz). To avoid phase discontinuity, the broadband tagging signal (Fig. 1B) was generated using phase modulation as follows,

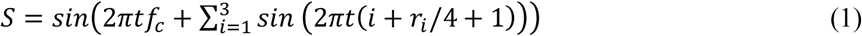

where *f*_*c*_ denotes frequency of the carrier signal (65 Hz), *t* is a time variable (0 – 2 s) and *r*_*i*_ is a random number from uniform distribution [0, 1]. The tagging signals for the left and right stimuli were generated independently for each trial, and these signals were uncorrelated in the 2 seconds interval, i.e., the correlation was below 0.1. Direction of attention, pairing of face-house stimuli and tagging frequencies were counterbalanced over trials.

#### Projector

We used the PROPixx DLP LED projector (VPixx Technologies Inc., Canada) to present the visual stimuli. This projector provides a refresh rate up to 1440 Hz by dividing each frame received from the graphics card (at 120 Hz) into multiple frames. The projector divides each received frame (1920 x 1200 pixels) into four equally sized quadrants (960 x 600 pixels), allowing for a fourfold increase in refresh rate (480 Hz). Colour (RGB) images presented in each quadrant are further converted to a grayscale representation by equalizing all components of RGB code. This allows for a 1440 Hz refresh rate since the 120 Hz is multiplied by a factor of respectively 4 and 3 fold when presenting grayscale images with a resolution of 960 x 600 pixels.

#### MEG and MRI data acquisition

MEG was acquired using a 306-sensor TRIUX Elekta Neuromag system (Elekta, Finland). The MEG data were lowpass filtered at 300 Hz using embedded anti-aliasing filters and sampled at 1000 Hz. Head position of the participants was digitized using the Polhemus Fastrack electromagnetic digitiser system (Polhemus Inc., USA). We also used an EyeLink eye tracker, and vertical and horizontal EOG sensors to remove trials containing blinks and saccades.

The tagging signals were recorded using a custom-made photodetector (Aalto NeuroImaging Centre, Aalto University, Finland) that was connected to a miscellaneous channel of MEG system. This allowed us to acquire the tagging signal with the same temporal precision as the MEG data.

A high-resolution T1-weighted anatomical image (TR/TE of 7.4/3.5 ms, a flip angle of 7°, FOV of 256×256×176 mm, 176 sagittal slices, and a voxel size of 1×1×1 mm^3^) was acquired using 3-Tesla Phillips Achieva scanner.

#### MEG data preprocessing

MEG data were analysed using MATLAB and the Fieldtrip toolbox (Oostenveld et al., 2011). The data were segmented into 3.5 s epochs; -1.0 – 2.5 s relative to the onset of the flickering stimuli (houses and faces).

Eye blinks were detected in the X-axis and Y-axis channels of the eye-tracker by applying a threshold of 5 SD. The saccades were detected using scatter diagram (or joint histogram) of X-axis and Y-axis time series of the eye-tracker for each trial. An event was classified as a saccade if the focus away from the central cross (i.e. fixation point) lasted longer than 500 ms. The trials contaminated by blinks and saccades were removed from further analysis, and the total amount of such trials was not exceeded 10%.

#### Attention modulation index

We quantified the effect on attention on the power at the alpha power and tagging signal using attention modulation index (AMI). To this end, spectral power was computed using Fourier transform (FT) for each MEG sensor and each epoch from 0 to 2 s relatively to stimulus onset:

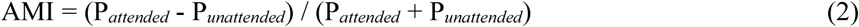

where P_*attended*_ and P_*unattended*_ denote spectral power averaged over attended and unattended stimuli trials, respectively.

For correlation analysis, we calculated individual AMI in the alpha band (8–13 Hz) for left and right sensors separately and then combined AMI over the sensors (see, Zhigalov et al., 2019).

### Sensor space data analysis

To assess coupling strength and latencies between the tagging signal and neuronal response, we computed the phase-locking value (*PLV*; Lachaux et al., 1999). The signals were filtered using 4^th^ order Butterworth zero-phase filter implemented as combination of high-pass and low-pass filters with cut-off frequencies 55 Hz and 75 Hz, respectively, and PLV was computed as follows,

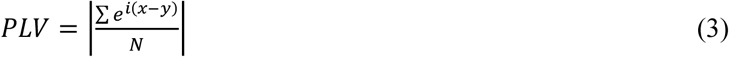

where *x* and *y* are instantaneous phases of the filtered signals (i.e. MEG and tagging signal), *N* is the number of samples and ‖ denotes an absolute value. The instantaneous phase was estimated using Hilbert transform after bandpass filtering. The bandwidth of the filter was slightly larger (55 – 75 Hz) than the bandwidth of the tagging signal (60 – 70 Hz) to avoid aliasing.

The phase-locking values were computed between the tagging signal and MEG signal (0 – 2 s from stimulus onset, Fig. 2A) at multiple lags. This is akin to calculating the cross-correlation, where the correlation coefficient is replaced by PLV. These cross-PLVs were estimated for each condition, and then maximum value of PLV and corresponding latency were assessed for each MEG sensor and condition. The cross-PLV and latencies were averaged separately for trials comprising attended and unattended stimuli (Fig. 2B).

**Figure 2.**
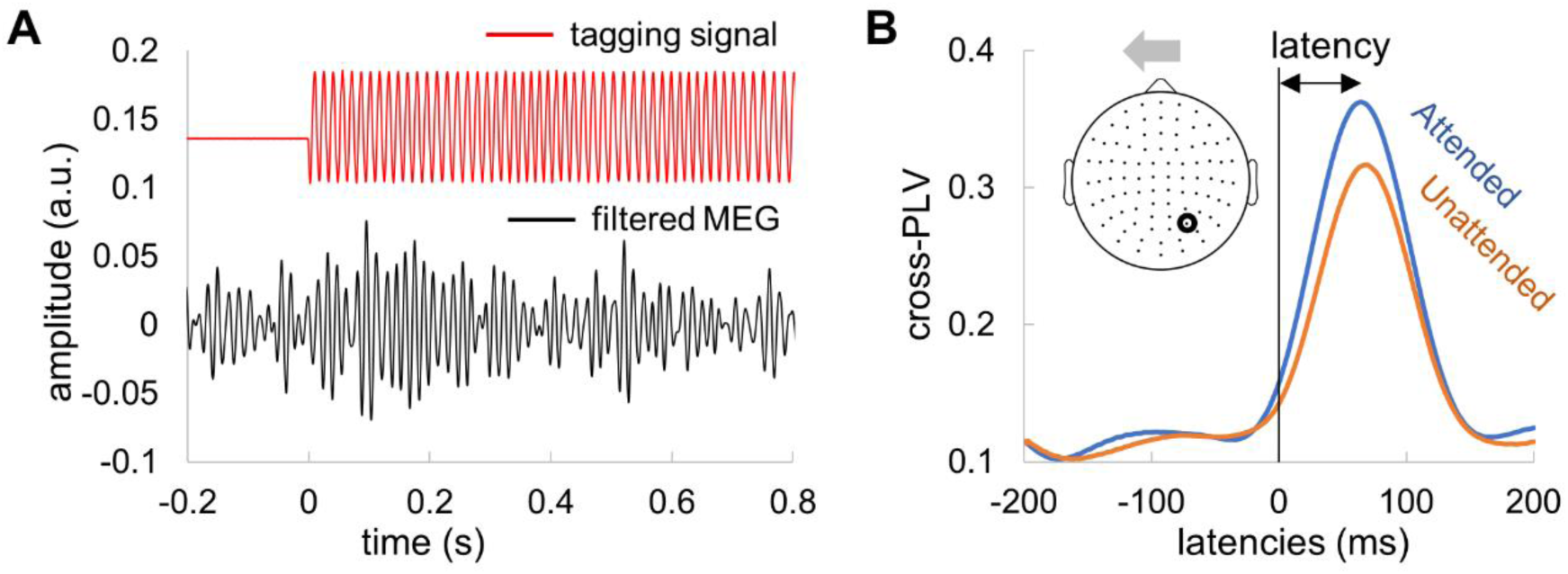
Estimation of response latency using the broadband frequency tagging technique. (A) Example of a single trial tagging visual signal (recorded by photodetector attached to the projector screen) and the MEG response. (B) An example of cross-PLV at multiple lags. The signals for attended compared to unattended objects showed larger cross-PLV. The peak of the cross-PLV reflects the delay in the MEG signals with respect to the visual input drive.

### Source space data analysis

AMI was computed in source space using dynamical imaging of coherent sources (DICS; (Gross et al., 2001)) spatial filter approach as implemented in Fieldtrip (Oostenveld et al., 2011).

To build a forward model, we first manually aligned the MRI images to the head shape digitization points acquired with the Polhemus Fastrak. Then, the MRI images were segmented, and a single shell head model was prepared using surface spherical harmonics fitting the brain surface (Nolte, 2003).

The time-frequency analysis (using multi-taper frequency transformation as implemented in Fieldtrip) has been applied to the data in the interval 0 – 2 s after flickering stimuli onset. DICS spatial filters were computed for 10 mm grid where individual anatomy was warped into standard MNI template. Finally, AMI (see, equation 1) was computed for each grid point at the alpha frequency (8 – 13 Hz) and the tagging response (55 – 75 Hz).

## Results

In this study, participants performed a cued visual attention task in which stimuli were frequency tagged using independent 60–70 Hz broadband signals in the left and right visual hemifield.

### Power of neuronal response modulated by spatial attention

We first quantified the modulations of neuronal activity by calculating the time-frequency representation of power. Next, we calculated the attention modulation index (AMI) be subtracting the alpha power for *attended* from *unattended* trials in the cue-target interval (0 – 2 s). The group level AMI was well in line with our previous observations for the alpha power and the tagging responses at 55–75 Hz (Fig. 3A). The alpha power decreased contra-laterally (and increased ipsi-laterally) to the attended hemifield (∼10%). An opposite pattern was observed at the tagging responses, where power of the tagging response increased contra-laterally (and decreased ipsi-laterally) to the attended hemi-field (∼5%). We applied a source modelling based on a beamforming approach to estimate the locations of the underlying neuronal sources. While the alpha power was localized in the parieto-occipital cortex, the high frequency tagging response was generated in the primary visual cortex (Fig. 3B).

**Figure 3.**
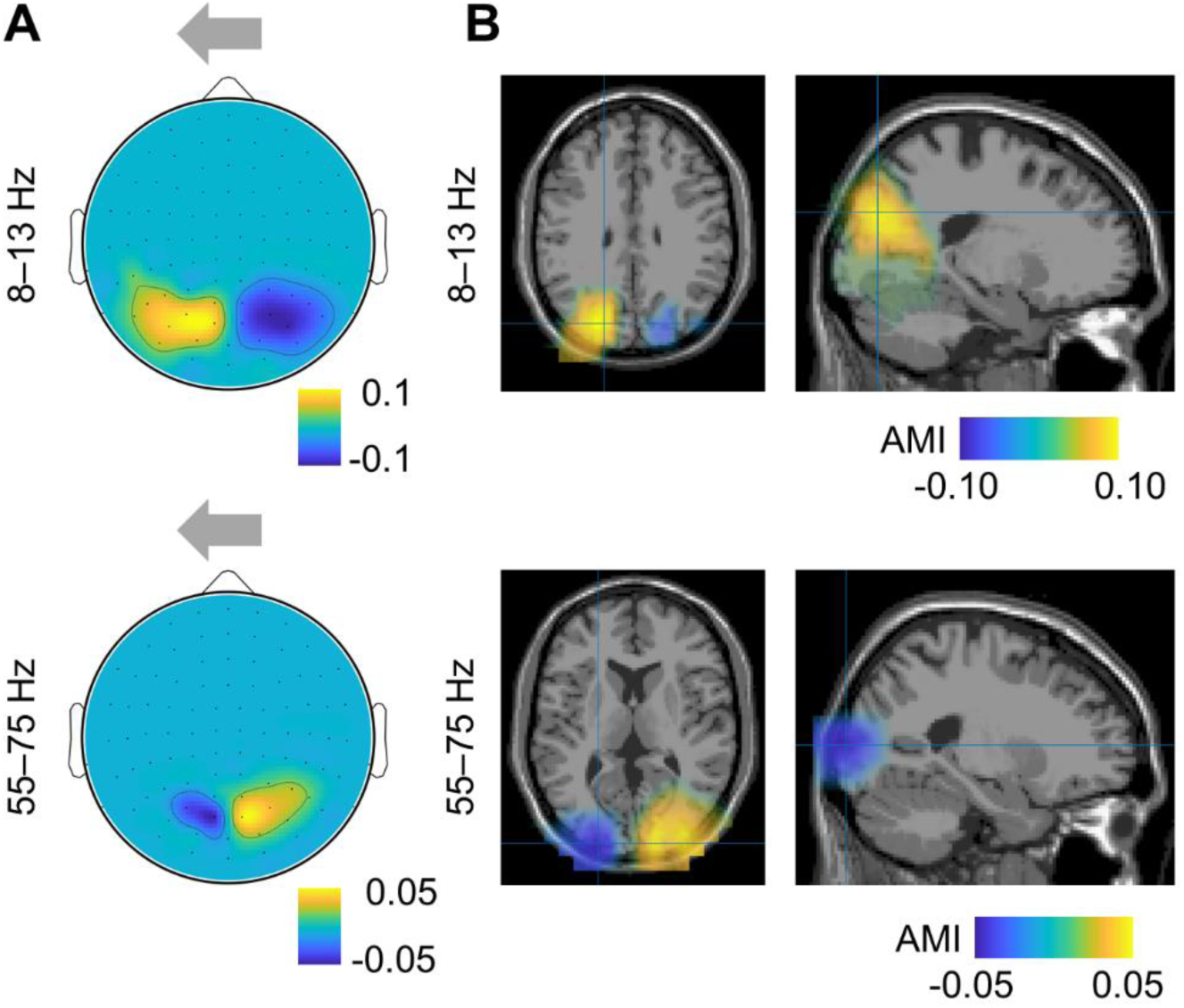
Attention modulates power of the alpha oscillations (8 – 13 Hz) and tagging response (55 – 75 Hz). The attention modulation index (AMI) was calculated by subtracting the estimated power for unattended (here, *attend right*) from attended (here, *attend left*) trials in the cue target interval (0 – 2 s). (A) Sensor-level AMI for the alpha oscillations (top) and tagging response (bottom). (B) Source estimates of the AMI for the alpha (top) and tagging response (bottom). Note the sources around the parieto-occipital sulcus for the alpha band modulation and early visual cortex for the tagging response.

### Sources of alpha and tagging response are spatially distinct

It can be readily seen in both sensor- and source-level analyses (Fig. 2) that the spatial patterns for the alpha power and the tagging responses are distinct. Next, we assessed the individual distances between the source peaks of the alpha and frequency tagged response (Fig. 4A). The yellow and blue markers indicate respectively the individual locations of the strongest alpha and frequency tagging modulations. Statistical analysis showed that the alpha modulation was on average about 5 mm superior to the modulation of the tagging responses (*p* < 0.001, pairwise t-test; Fig. 4B). The effect was similar for both left and right hemispheres. While the sources of the tagging response were largely located in the primary visual cortex, sources of the alpha oscillations were located closer to posterior parietal cortex. Similar observations but without rigorous testing have been made in a spatial attention paradigm in the alpha and gamma frequencies (Bauer et al., 2014). Our results suggest that sources of the alpha and tagging responses are spatially distinct and therefore likely to support different mechanisms.

**Figure 4.**
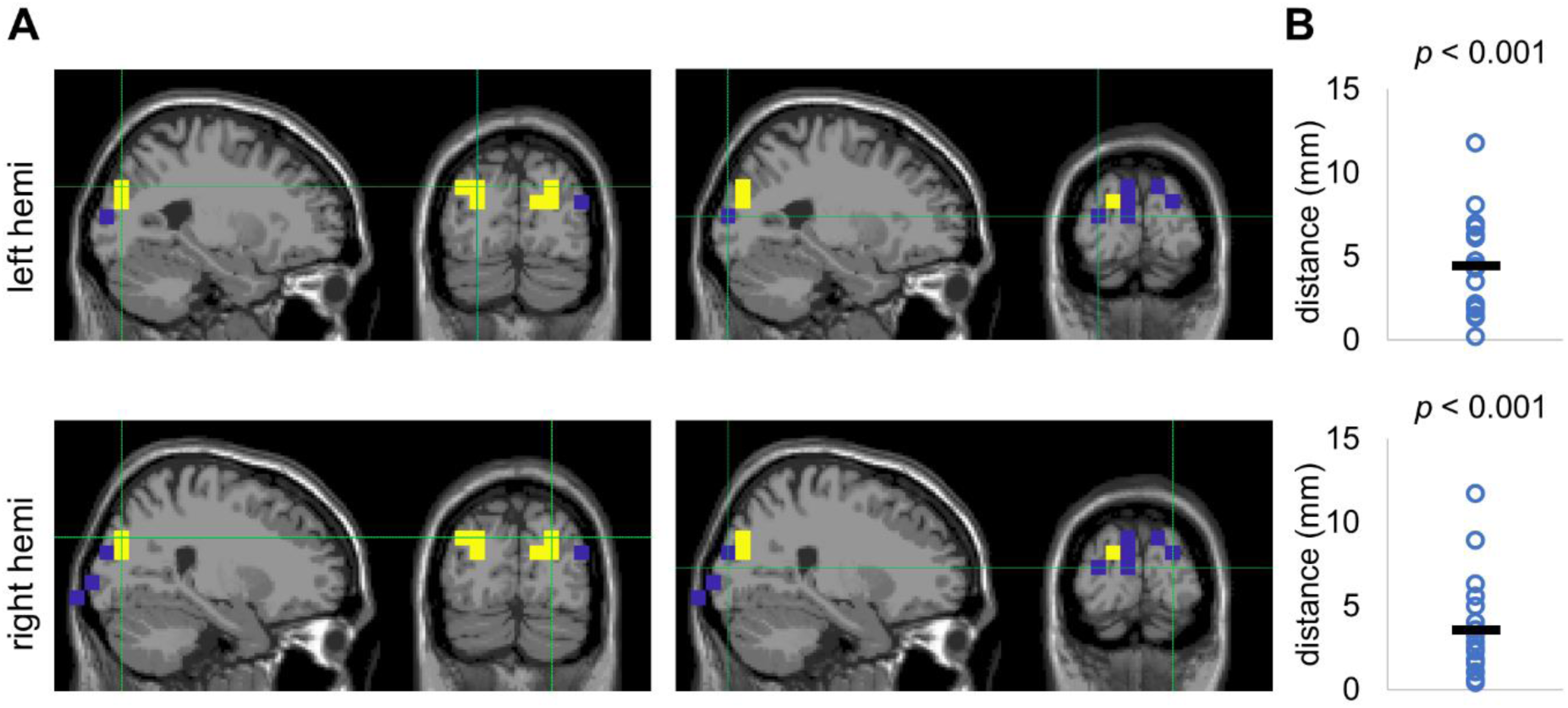
Source-level AMI for the alpha and tagging response are spatially distinct. (A) The peak individual values of alpha AMI (yellow dots; left plots) were located around the parieto-occipital cortex while largest individual values of tagging AMI (blue dots; right plots) were located closer to the primary visual cortex. (B) The distances between the peak locations of the AMI for the alpha and tagging response in individual subjects were significantly larger than zero in both hemispheres.

### Alpha power is not related to the power of tagging response

To further investigate the relationship between the alpha power and tagging response modulation, we computed – per subject – the power of the tagging response with respect to the median split of the trials according to alpha power (Fig. 5A). The frequency tagging responses with respect to low and high alpha power were the averaged over subjects and then compared using the pairwise t-test (Fig. 5B). Pairwise comparison revealed a complete absence of a relation (*p* > 0.58, pairwise t-test). These results support the earlier notion that although the power of the alpha and tagging response are modulated by attention, they are not directly coupled.

**Figure 5.**
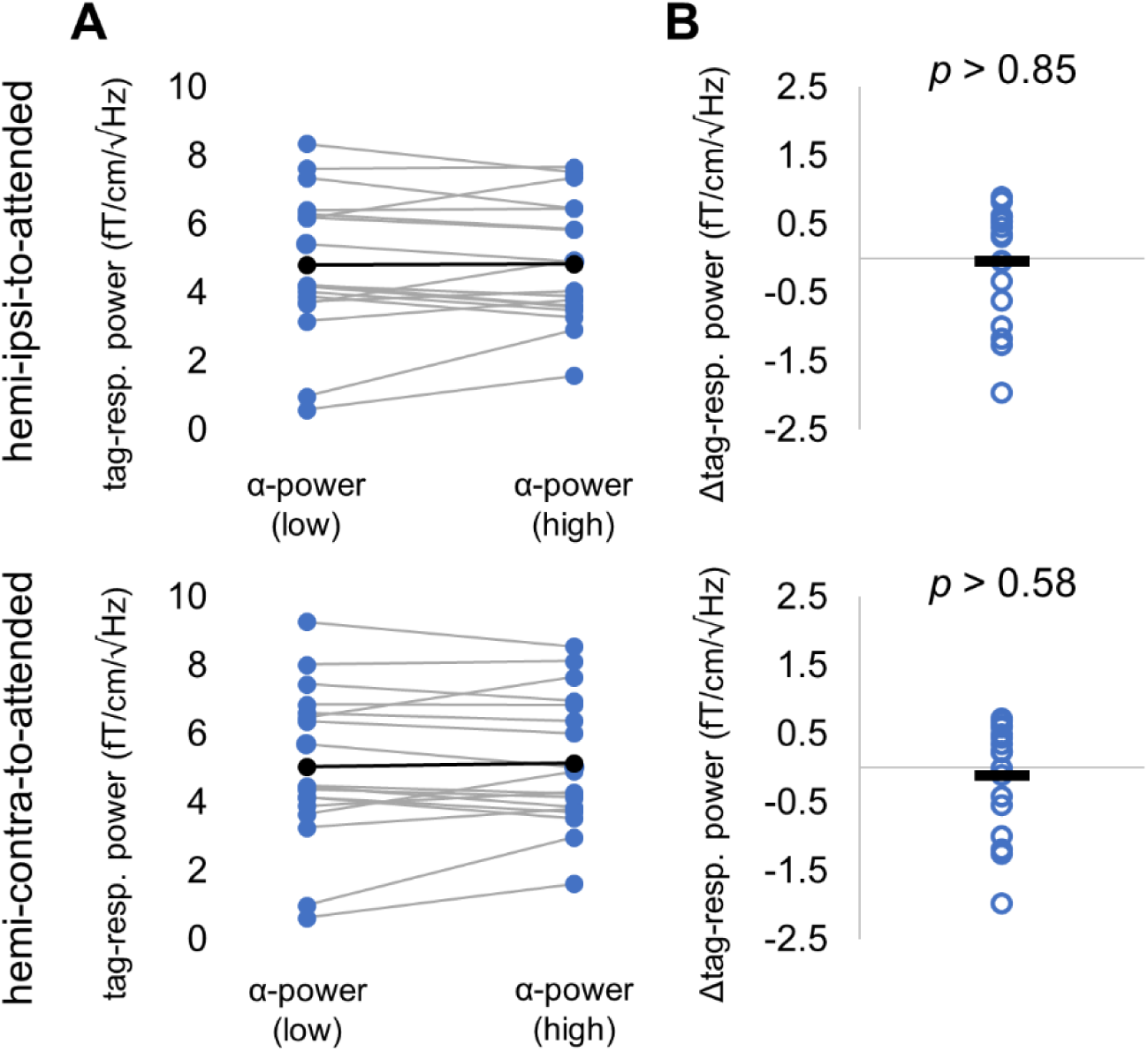
The tagging response power with respect to the median split of low and high alpha power over trials. (A) Power of the tagging response (denoted as tag-resp. power) for individual subjects when pooled over trials with respect to low and high alpha power. (B) Pair-wise difference of the data shown in A. These findings demonstrate an absence in correlation between the alpha power and the frequency tagged signals.

### Attention modulates increased the magnitude of the tagged MEG response

Similarly to the power of tagging response at 55–75 Hz (Fig. 3A), the cross-PLVs at lags 45 – 60 ms (see methods, Fig. 2B) were larger for attended compared to unattended stimuli in the occipital sensors. To assess the difference statistically, we selected the strongest responding sensor (planar gradiometers) when considering the combined conditions for each subject. We observed significant differences in the cross-PLV measure between tagging and MEG signals (Fig. 6) related to attention (*p* < 0.001 for left and as well as right sensors of interest, paired t-test). These results are in line with earlier findings (Zhigalov et al., 2019) as well as Figure 3 showing that attention increases the magnitude of the tagged response.

**Figure 6.**
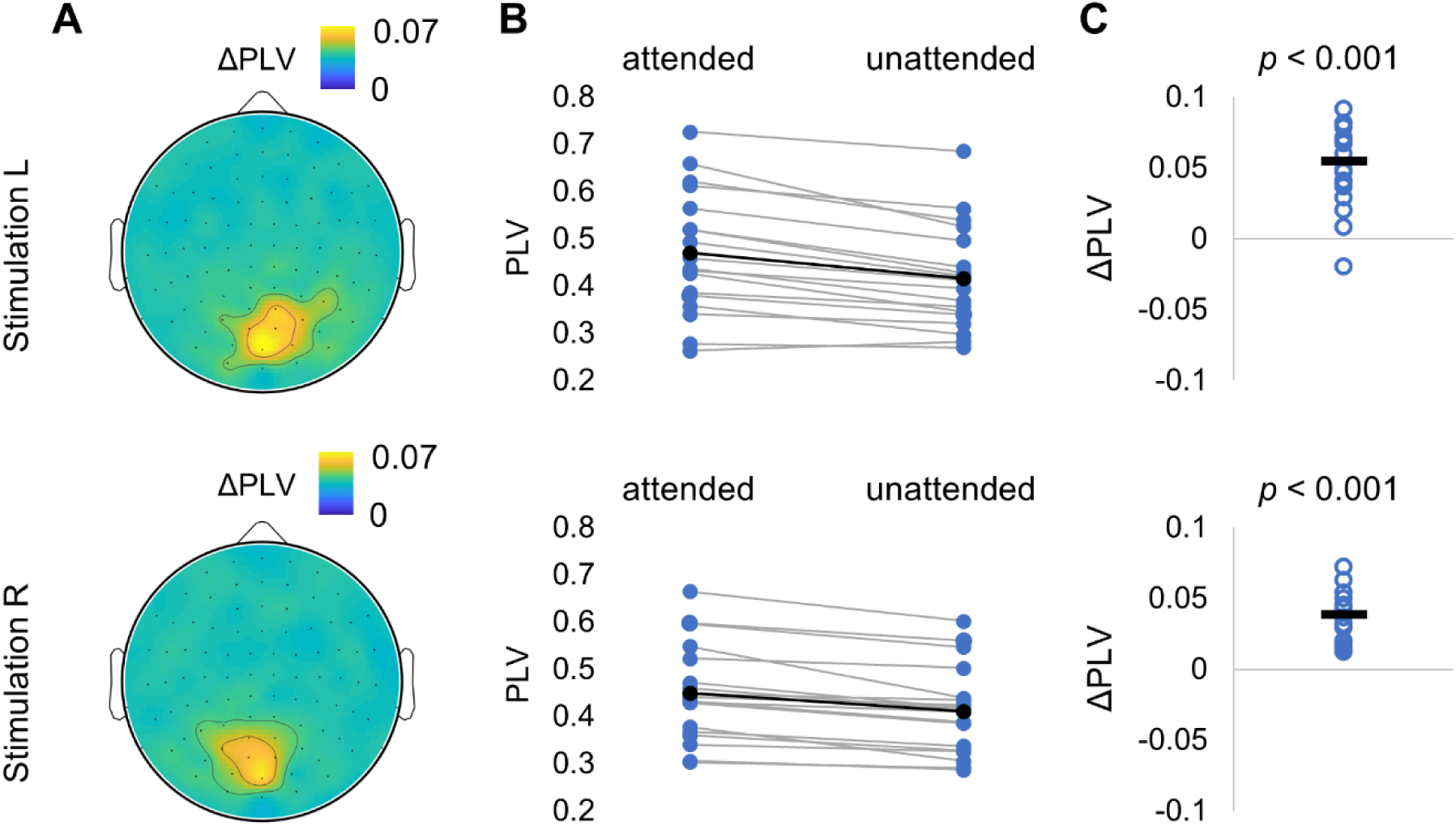
Attention modulates coupling strength between the tagging signal and MEG. (A) Spatial patterns (topographies) of the cross-phase-locking values (cross-PLV) at 45 – 60 ms between the visual drive (photo diode) and the MEG signal at group level for attended and unattended stimuli for the stimulation on the left (top panel) and on the right (bottom panel). The cross-PLV was derived in the 0 – 2 s interval after stimulation onset and was averaged over planar gradiometers. (B, C) Pairwise comparison showed a significant increase in cross-PLV (lag 45 – 60 ms) for attended versus unattended objects.

### Attention does not affect latency of the tagging MEG response

To assess the latency between the visual input and the MEG response, we estimated the lag of the maximum cross-PLV (see methods, Fig. 2B). We found that attention did not affect the latency between the neuronal response and the tagging signal in the occipital sensors of interest (Fig. 7A). For statistical comparison, we considered the strongest responding sensor left and right sensors for each subject, in the same manner as it has been done for PLV (Fig. 6B). The results did not show a robust differences in neuronal response latencies with attention (Fig. 7B,C).

**Figure 7.**
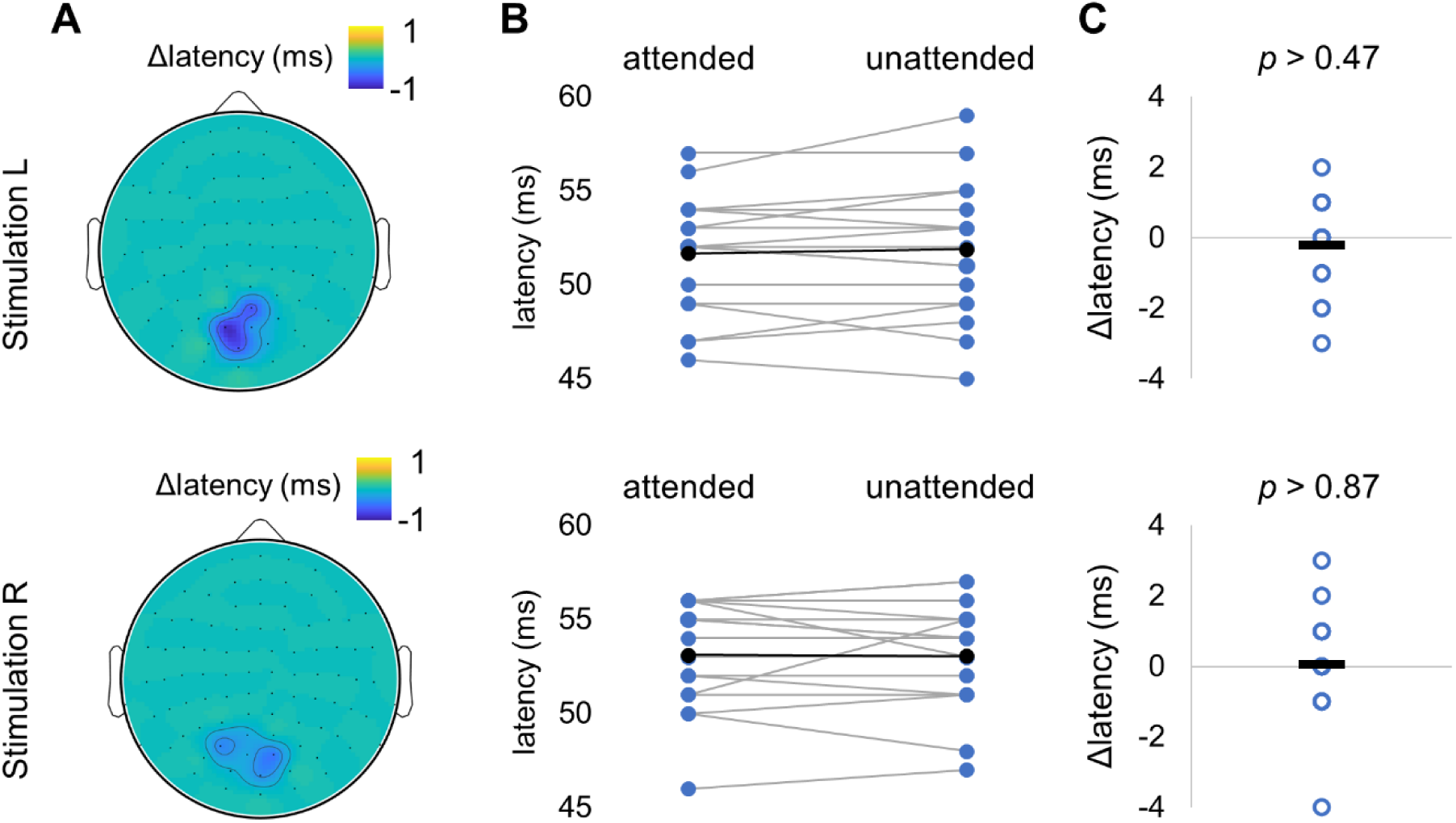
Attention did not affect the neuronal response latency between tagging signal and MEG. (A) Group level spatial patterns of latencies for attended and unattended stimuli for the stimulation on the left (top panel) and on the right (bottom panel). (B, C) Pairwise comparison did not show any significant changes in latencies for attended versus unattended objects.

### Relationship between power of ongoing alpha activity and parameters of tagging response

We assessed the relationship between individual AMI at the alpha band and AMI of tagging response, and also between the individual AMI at the alpha band and response latencies. First, we computed the correlation between individual AMI at the alpha frequency and tagging response at 55-75 Hz (Fig. 8A). These results showed a negative correlation (*r* = -0.63 (Spearman correlation), *p* < 0.005) between the modulation of power at the alpha and tagging responses, which is in line with earlier observations (Zhigalov et al., 2019). Similarly, we correlated individual AMI in the alpha band and the response latencies (Fig. 8B); however, the correlation was not significant (*r* = -0.20 (Spearman correlation), *p* > 0.43).

**Figure 8.**
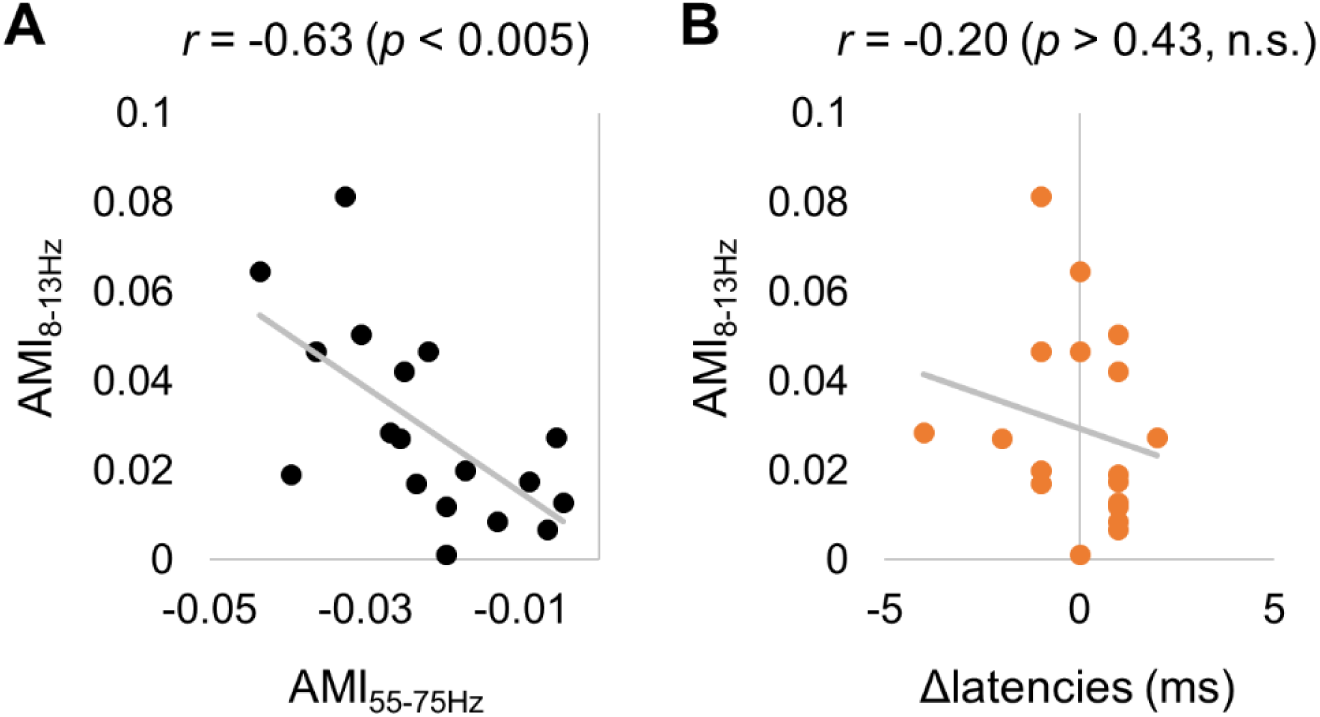
Relationship between attentional modulation of the alpha power and the tagging response over subjects. (A) The decrease in alpha power with attention was correlated with an increase in the tagging response with attention over subjects. (B) Attentional modulation of the alpha power was not related to attentional modulation of the response latency over subjects.

## Discussion

In this MEG study we used a spatial attention task in combination with a novel broadband frequency tagging technique. We found that the power of alpha activity and the tagging response were modulated by attention, confirming earlier results (Zhigalov et al., 2019). Source modelling of the MEG data allowed us to identify the neuronal generators of the frequency tagging signal in early visual regions, and the alpha oscillations generators around parieto-occipital sulcus. Importantly we showed that the power of alpha and tagging response were not related at single trial level. By further analysing the broadband tagging response we showed that the response delays were not modulated by attention.

### Neuronal excitability and alpha change with attention

Numerous studies have demonstrated that alpha oscillations are top-down modulated when attention is allocated (Foxe and Snyder, 2011; Klimesch, 2012; Müller and Weisz, 2012). This has resulted in the idea that alpha oscillations serve to control neuronal gain in early visual regions (Jensen and Mazaheri, 2010; Spaak et al., 2012; Jensen et al., 2015). Using multi-contact laminar electrodes to measure spontaneous signals simultaneously from all layers of V1, Spaak and colleagues (Spaak et al., 2012) found a robust coupling between alpha phase in the deeper layers and gamma amplitude in granular and superficial layers. In the same vein, a study by (Jensen et al., 2015) proposed layer- and frequency-specific mechanism of feedforward and feedback visual processing, in which alpha oscillations could modulate gamma activity in V1 area. We here operate under the premise that neuronal excitability and gain control can be quantified by means of the frequency tagged response.

Our findings challenge the notion that alpha oscillations exert gain control in early sensory regions. First, the power at the alpha and tagging responses were not correlated at the single trial level as also demonstrated in previous findings (Zhigalov et al., 2019; Gundlach et al., 2020). Second, the sources of the alpha oscillations were localized around the parieto-occipital sulcus while the sources of tagging response were located in primary visual cortex. We conclude that alpha oscillations do not serve to adjust the gain in early visual regions; rather the alpha oscillations serve to gate neuronal activity in regions downstream of the visual cortex. As shown in Figure 9, we propose that this gating serves the allocation of neuro-computational resources and modulates the feedforward flow in parieto-occipital areas. As proposed in previous work (Jensen and Mazaheri, 2010), the gating could be implemented neurophysiologically by GABAergic pulsed inhibition in the alpha frequency range that will limit neuronal processing and the feedforward flow.

**Figure 9.**
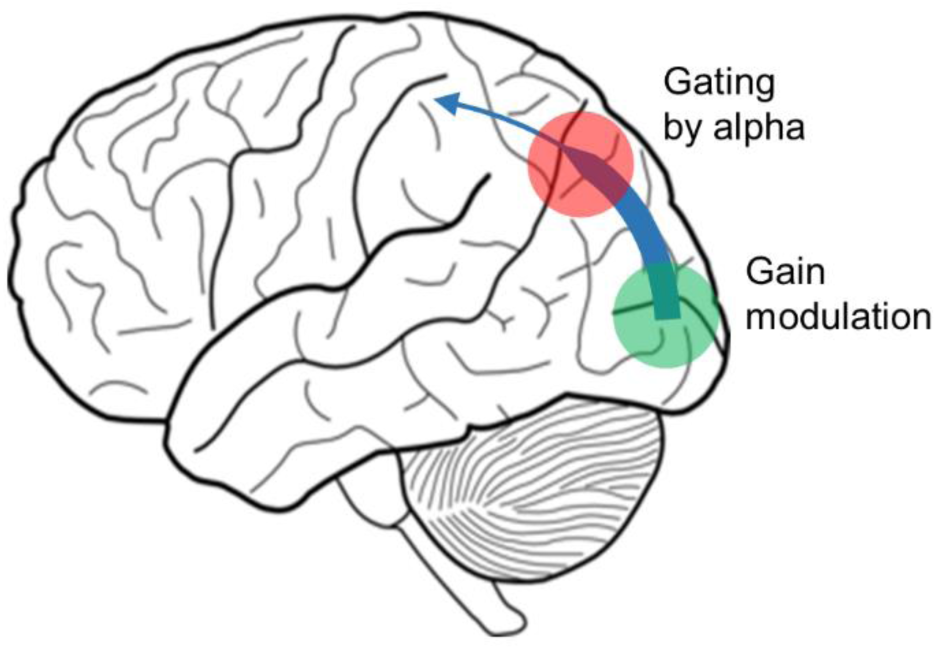
We find that spatial attention modulates the frequency tagging response in early visual regions (green) and alpha oscillations around the parieto-occipital sulcus (red). Importantly the alpha activity and the frequency tagging response were not correlated over trials. This points to a scheme in which alpha oscillations gate the information flow in down-stream visual regions without directly controlling the gain in early visual regions.

### Attention does not modulate latencies of the tagging response

It is debated to what extend attention modulates the speed of visual processing. Several electrophysiological studies in monkeys (Lee et al., 2007; Lee and Maunsell, 2010) showed that contrast but not attention modulated the latency of the response. However, recent studies (Sundberg et al., 2012; Galashan et al., 2013) showed that attention impacted the latencies of neuronal responses in V4 and MT areas respectively, although the effect was relatively small (1 – 2 ms). Using broadband frequency tagging, we estimated the delay in neuronal activation with respect to the visual input by time-shifting the two signals in order to identify when they were strongest coupled. Consistent with the literature (e.g., (Lee and Maunsell, 2010; Maunsell, 2015)) we found that the visual cortex responded maximally ∼50 ms after the visual input; however, attention did not modulate the response latencies. Consistent with prevailing views (Tallon-Baudry, 2012) we conclude that spatial attention does not modulated response latencies in early visual cortex; however, our approach does provide an exciting new tool for estimating the delay of neuronal activation in visual cortex.

### Power of the response but not latency of the tagging response is affected by ongoing alpha activity at group level

We replicated earlier observations demonstrating that attentional modulation of power of the alpha and tagging responses are negatively related at group level (Zhigalov et al., 2019). However, there was no evidence that the alpha power was correlated with the response latencies. The over-subject correlation does point to a relation between neuronal excitability in early visual regions and alpha power; however, this effect might be partly explained by different signal-to-noise ratios in the participants and thus not functional.

### Conclusion

We conclude that alpha oscillations are not directly involved in gain control of neuronal activity in early visual regions. Rather the alpha oscillations might serve to gate the feed-forward flow in downstream regions e.g. around the parieto-occipital sulcus. In future work it would be of great interest to uncover the neuronal mechanisms implementing the gating in relation to the alpha oscillations as well as the gain control as reflected by the rapid frequency tagging responses.

## Acknowledgment

The work was supported by the following funding: a James S. McDonnell Foundation Understanding Human Cognition Collaborative Award (grant number 220020448), the Wellcome Trust Investigator Award in Science (grant number 207550), a BBSRC grant (BB/R018723/1) as well as the Royal Society Wolfson Research Merit Award.

## References

Bauer M, Stenner MP, Friston KJ, Dolan RJ (2014) Attentional modulation of alpha/beta and gamma oscillations reflect functionally distinct processes. Journal of Neuroscience 34:16117–16125.

Desimone R, Duncan J (1995) Neural Mechanisms of Selective Visual Attention. Annual Review of Neuroscience 18:193–222.

Foxe JJ, Snyder AC (2011) The Role of Alpha-Band Brain Oscillations as a Sensory Suppression Mechanism during Selective Attention. Frontiers in Psychology 2:154.

Galashan FO, Saßen HC, Kreiter AK, Wegener D (2013) Monkey area MT latencies to speed changes depend on attention and correlate with behavioral reaction times. Neuron 78:740–750.

Gross J, Kujala J, Hamalainen M, Timmermann L, Schnitzler A, Salmelin R (2001) Dynamic imaging of coherent sources: Studying neural interactions in the human brain. Proceedings of the National Academy of Sciences of the United States of America 98:694–699.

Gulbinaite R, Roozendaal DHM, VanRullen R (2019) Attention differentially modulates the amplitude of resonance frequencies in the visual cortex. NeuroImage 203:116146.

Gundlach C, Moratti S, Forschack N, Müller MM (2020) Spatial Attentional Selection Modulates Early Visual Stimulus Processing Independently of Visual Alpha Modulations. Cerebral cortex (New York, NY : 1991).

Herbst SK, Javadi AH, van der Meer E, Busch NA (2013) How Long Depends on How Fast— Perceived Flicker Dilates Subjective Duration Meck WH, ed. PLoS ONE 8:e76074.

Jensen O, Bonnefond M, Marshall TR, Tiesinga P (2015) Oscillatory mechanisms of feedforward and feedback visual processing. Trends in Neurosciences 38:192–194.

Jensen O, Hanslmayr S (2020) The Role of Alpha Oscillations for Attention and Working Memory (David Poeppel, George Mangun MG, ed)., Sixth Edition. MIT Press.

Jensen O, Mazaheri A (2010) Shaping functional architecture by oscillatory alpha activity: Gating by inhibition. Frontiers in Human Neuroscience 4.

Kastner S, Nobre A (2014) The Oxford Handbook of Attention.

Keitel C, Keitel A, Benwell CSY, Daube C, Thut G, Gross J (2019) Stimulus-driven brain rhythms within the alpha band: The attentional-modulation conundrum. Journal of Neuroscience 39:3119–3129.

Kleiner M, Brainard D, Pelli D, Ingling A, Murray R, Broussard C (2007) What’s new in psychtoolbox-3. [Pion Ltd.].

Klimesch W (2012) Alpha-band oscillations, attention, and controlled access to stored information. Trends in Cognitive Sciences 16:606–617.

Lee J, Maunsell JHR (2010) Attentional modulation of MT neurons with single or multiple stimuli in their receptive fields. Journal of Neuroscience 30:3058–3066.

Lee J, Williford T, Maunsell JHR (2007) Spatial attention and the latency of neuronal responses in macaque area V4. Journal of Neuroscience 27:9632–9637.

Mathewson KE, Lleras A, Beck DM, Fabiani M, Ro T, Gratton G (2011) Pulsed Out of Awareness: EEG Alpha Oscillations Represent a Pulsed-Inhibition of Ongoing Cortical Processing. Frontiers in Psychology 2:99.

Maunsell JHR (2015) Neuronal Mechanisms of Visual Attention. Annual Review of Vision Science 1:373–391.

Müller N, Weisz N (2012) Lateralized auditory cortical alpha band activity and interregional connectivity pattern reflect anticipation of target sounds. Cerebral cortex (New York, NY : 1991) 22:1604–1613.

Nolte G (2003) The magnetic lead field theorem in the quasi-static approximation and its use for magnetoenchephalography forward calculation in realistic volume conductors. Physics in Medicine and Biology 48:3637–3652.

Oostenveld R, Fries P, Maris E, Schoffelen J-M (2011) FieldTrip: Open source software for advanced analysis of MEG, EEG, and invasive electrophysiological data. Computational intelligence and neuroscience 2011:156869.

Romei V, Brodbeck V, Michel C, Amedi A, Pascual-Leone A, Thut G (2008) Spontaneous fluctuations in posterior α-band EEG activity reflect variability in excitability of human visual areas. Cerebral Cortex 18:2010–2018.

Spaak E, Bonnefond M, Maier A, Leopold DA, Jensen O (2012) Layer-specific entrainment of γ-band neural activity by the α rhythm in monkey visual cortex. Current biology : CB 22:2313–2318.

Sundberg KA, Mitchell JF, Gawne TJ, Reynolds JH (2012) Attention influences single unit and local field potential response latencies in visual cortical area V4. Journal of Neuroscience 32:16040–16050.

Tallon-Baudry C (2012) On the neural mechanisms subserving consciousness and attention. Frontiers in Psychology 3.

van Diepen RM, Foxe JJ, Mazaheri A (2019) The functional role of alpha-band activity in attentional processing: the current zeitgeist and future outlook. Current Opinion in Psychology 29:229–238.

Watson AB, Pelli DG (1983) Quest: A Bayesian adaptive psychometric method. Perception & Psychophysics 33:113–120.

Willenbockel V, Sadr J, Fiset D, Horne GO, Gosselin F, Tanaka JW (2010) Controlling low-level image properties: The SHINE toolbox. Behavior Research Methods 42:671–684.

Zhigalov A, Herring JD, Herpers J, Bergmann TO, Jensen O (2019) Probing cortical excitability using rapid frequency tagging. NeuroImage 195:59–66.

